# AmpC β-lactamases: A Key to Antibiotic Resistance in ESKAPE Pathogens

**DOI:** 10.1101/2025.03.05.641667

**Authors:** Deeksha Pandey, Isha Gupta, Dinesh Gupta

## Abstract

AmpC β-lactamases (*blaAmpC*) are one of the important drivers for a high incidence of antimicrobial resistance (AMR) within ESKAPE pathogens, which are a group of bacteria that are the leading cause of hospital-acquired infections. Investigating the presence and characteristics of the *blaAmpC* is essential for understanding the molecular mechanism of resistance and developing effective strategies to combat AMR. We have investigated the presence of *blaAmpC*, distribution on plasmids/chromosomes, copy number, amino acid identities/variabilities, and evolutionary characteristics in ESKAPE pathogens. Our analysis indicates 1790 AmpC enzymes in 4713 completely assembled genomes, subdivided into nine different enzyme groups. AmpC is more common in *A. baumannii*, followed by *P. aeruginosa, K. pneumoniae, and Enterobacter spp*., but absent in *E. faecium* and *S. aureus.* Among the nine AmpC enzyme groups, ACC, ACT, CMH, and MIR are exclusively found in *Enterobacter spp.*, while the largest enzyme group ADC is only present in *A. baumannii*; PDC and PIB are found only in *P. aeruginosa*. Phylogenetic analysis revealed significant divergence amongst few enzyme groups and closer evolutionary relationships within others. Functional motif analysis identified conserved catalytic residues across all enzyme groups, except PIB, which demonstrates structural and functional divergence. Because of these variations, PIB’s ability to bind cephalosporins decreases while enhancing its activity against carbapenems, hence, we have excluded PIB from further analysis. This is the first detailed report on AmpC in ESKAPE pathogens, which also re-emphasizes the wise use of antibiotics to reduce the rise of AmpC cephalosporins resistance, a significant public health concern.

**Highlights:**

1. First detailed report on ESKAPE *blaAmpC* localization in chromosomes and plasmids
2. *blaAmpC* is absent in *E. faecium* & *S. aureus* but prevalent in other four pathogens
3. Most *blaAmpC* belong to ADC, exclusively present in *A. baumannii* chromosomes
4. Interspecies gene exchange in *K. pneumoniae*, *P. aeruginosa* & *Enterobacter spp*.
5. Evolutionary analysis shows recent divergence & possible horizontal gene transfer

## Introduction

Antimicrobial Resistance (AMR) is ranked among the top ten threats to global public health. The emergence and spread of multidrug-resistant (MDR) and extensively drug-resistant (XDR) bacteria, particularly *Enterobacteriaceae* and *Acinetobacter*, which are on the World Health Organization bacterial priority pathogens 2024 list named as ESKAPE (*Enterococcus faecium*, *Staphylococcus aureus*, *Klebsiella pneumoniae*, *Acinetobacter baumannii*, *Pseudomonas aeruginosa*, and *Enterobacter spp.*), pose a serious threat to human and animal health. MDR bacterial infections are a serious public health concern, accounting for a significant share of morbidity and mortality worldwide [1, 2]. The global proliferation of MDR bacteria has led to a heightened incidence of in-hospital mortality and a scarcity of treatment options, rendering it an increasingly alarming issue. Bacterial AMR, responsible for around 4.95 million fatalities among them, especially ESKAPE pathogens, which caused 1.2 million deaths. ESKAPE pathogens are notorious bacteria that cause severe nosocomial infections worldwide. Cases of bacterial AMR patients are extremely difficult to treat and pose a serious threat to global public health due to their high ability to acquire antibiotic resistance [3, 4].

ESKAPE pathogens develop resistance through diverse genetic mechanisms, including point mutations, amplification of genomic segments, and horizontal transfer of resistance genes [5]. ESKAPE pathogens employ multiple approaches to neutralize the antibiotics’ effect which include (i) enzymatic degradation, (ii) efflux pumps, (iii) target modification and (iv) biofilm formation [6]. Among the four major bacterial antibiotic resistance mechanisms β-lactamase production stands out as one of the most clinically significant. This is because β-lactam antibiotics (penicillins, cephalosporins, monobactams, and carbapenems) form the cornerstone of modern antimicrobial therapy [7]. The production of β-lactamases by bacteria is a critical factor in their ability to evade these antibiotics, posing a significant threat to global public health. Some β-lactamases are encoded on mobile genetic elements (such as plasmids), while others are encoded on chromosomes. Penicillins and other β-lactam antibiotics have served as the backbone of modern medicine, affecting a range of bacterial infections [8]. The key to this problem appears to be found in the enzymes that bacteria produce that inactivate these life-saving drugs; the β-lactamases [9]. β-lactamase, are classified by the Ambler molecular system into four classes: Class A (e.g., TEM, SHV, KPC), Class B (metallo-beta-lactamases like NDM), Class C (AmpC beta-lactamases (*blaAmpC*)), and Class D (oxacillinases like OXA) [7]. Of all the types of β-lactamases, the AmpC enzymes are of great concern as they can hydrolyze many different types of β-lactams including cephalosporins and penicillin, along with various inhibitors designed to counter β-lactamase activity [10]. *blaAmpC*, which are either chromosomally encoded or plasmid-mediated, pose a significant clinical threat due to their inducible expression, widespread dissemination (via plasmid transfer), and resistance to first- and second-generation cephalosporins. The emergence of *blaAmpC* producing bacteria has been established as an important public health concern, resulting in the need for rapidly evolving harmful strains that can produce resistance to all known treatments available. Understanding *blaAmpC* in detail is important for preventing the transmission of invasive organisms and improving infection control measures in clinical settings. Studying the phenomenon of *blaAmpC* production and evolution in such pathogens may shed light on their resistance mechanisms which is important in developing effective treatment and diagnostic measures [11]. To the best of our knowledge, this is the first detailed study to investigate the presence and diversity of *blaAmpC* enzyme groups, their amino acid identities, variabilities, and evolutionary characteristics among ESKAPE pathogens. This study aims to improve understanding of the resistive properties of *blaAmpC* and the mechanisms behind their transmission and contribute to the formulation of novel and better therapeutic and diagnostic strategies.

## 2. Results and Discussion

### 2.1. Data analysis and ESKAPE pathogen classification

The NCBI genome database consists of 3260, 15,739, 17,608, 8037, 8221, and 4970 entries, corresponding to *E. faecium, S. aureus, K. pneumoniae, A. baumannii, P. aeruginosa, and Enterobacter spp*., respectively. Out of these, 318, 1022, 1661, 579, 624, and 509 entries corresponded to complete genomes of *E. faecium, S. aureus, K. pneumoniae, A. baumannii, P. aeruginosa, and Enterobacter spp.,* respectively (**Table 1**). Among these complete genomes, *E. faecium* comprised 14 entries with only chromosomes and 304 belong to both chromosomes and plasmids. Similarly, *S. aureus* included 454 entries with only chromosomes and 568 with both chromosomal and plasmid. *K. pneumoniae* had 110 entries with only chromosomal DNA and 1551 entries with chromosomes and plasmids. *A. baumannii* comprised 122 entries with only chromosomes and 457 with both chromosome and plasmid. In *P. aeruginosa*, 497 entries corresponded to only chromosomes and 127 for both chromosome and plasmid. Furthermore, 98 entries corresponded to only chromosomes and 411 with both chromosome and plasmid in *Enterobacter spp*. The replicon-wise distribution analysis revealed that most ESKAPE pathogens showed plasmid(s) presence. The quantity of plasmids found in the ESKAPE pathogens showed an intriguing trend. The average number of plasmids per genome for *E. faecium, K. pneumoniae*, and *Enterobacter spp.* was roughly four. There were about two plasmids per genome in *A. baumannii*. However, the average number of plasmids per genome was one in the case of *S. aureus* and *P. aeruginosa*. Botelho *et al.* [12] also showed the same pattern that ESKAPE pathogens have a considerable fraction of plasmids encoding a significant percent of AMR, where over 35% of plasmids across the different ESKAPE species had at least one AMR gene.

**Table 1:** Replicon-Wise Genome Statistics and *blaAmpC* Distribution in ESKAPE Pathogens (NCBI Data) Summary of Genomic Data and *blaAmpC* Presence in ESKAPE Genomes from NCBI.

| Organism | Total no. of genomes (NCBI) | No. of complete genomes/no. of AmpC | Replicons |  |  |  |
| --- | --- | --- | --- | --- | --- | --- |
|  |  |  | No. of chromosomal genomes/no. of AmpC | Both Chromosome & Plasmid |  |  |
|  |  |  |  | No. of genomes containing both chromosomes & plasmids/ no. of AmpC (Total) | No. of chromosomes / no. of AmpC | No. of plasmids /no. of AmpC |
| *E. faecium* | 3,260 | 318/0 | 14/0 | 304/0 | 304/0 | 1420/0 |
| *S. aureus* | 15,739 | 1022/0 | 454/0 | 568/0 | 568/0 | 841/0 |
| *K. pneumoniae* | 17,608 | 1661/175 | 110/1 | 1551/11 | 1550/11 | 5594/152 |
| *A. baumannii* | 8,037 | 579/755 | 122/97 | 457/330 | 457/327 | 923/1 |
| *P. aeruginosa* | 8,221 | 624/401 | 497/250 | 127/75 | 127/75 | 188/1 |
| *Enterobacter* spp. | 4,970 | 509/459 | 98/19 | 411/207 | 411/233 | 1295/0 |
| **Total** | **57,835** | **4713/1790** | **1295/367** | **3418/623** | **3417/646** | **10261/154** |

### 2.2. Identification and filtration of *blaAmpC* sequences in ESKAPE pathogens

A profile HMM (pHMM) of *blaAmpC* was retrieved from the β-lacFampred tool, developed by us earlier [13]. pHMM profile of *blaAmpC* was searched against each of the six pathogens using the NHMMSCAN tool of the HMMER package [14]. Subsequent re-validation of the NHMMSCAN outputs against the GenBank annotations for the corresponding genomes did not identify any unannotated or additional AmpC. This indicates that the pHMM of *blaAmpC* showed 100% specificity for the AmpC family of class C β-lactamases. Using the Beta-Lactamase DataBase (BLDB) database, a specialized repository developed by Naas et al. [15], 3781 protein sequences of Class C β-lactamases were extracted for annotation of AmpC genes present in ESKAPE pathogens. A comprehensive BLAST search was performed across ESKAPE pathogen genomes against *blaAmpC* sequences. Sequences with less than 100% identity were eliminated, and duplicates were removed, resulting in a final dataset of 1790 unique *blaAmpC* sequences.

### 2.3. Dissemination and localization of *blaAmpC* in ESKAPE pathogens

1790 AmpC proteins were identified in the 4713 assembled ESKAPE pathogens genomes (**Table 1**). Interestingly, AmpC was completely absent in *E. faecium* and *S. aur*eus, but other pathogens showed a notable presence of the AmpC gene. The lack of AmpC in these two pathogens raises the possibility that their antibiotic resistance mechanisms are distinct.

*Enterobacter spp.* and *P. aeruginosa* genomes contained 459 and 401 AmpC sequences, respectively. AmpC was found on both chromosomes and plasmids in 75 *P. aeruginosa* and 207 Enterobacter *spp.* The average number of plasmids per genome varied among pathogens, with *K. pneumoniae* having the highest count, containing about 152 plasmids carrying AmpC sequences.

In *K. pneumoniae*, AmpC was identified in 1 out of 110 complete chromosomal genome assemblies, while 11 AmpC were located on chromosomes and 152 AmpC were discovered on plasmids among 1551 genomes containing both chromosomes and plasmids. This shows that AmpC is more frequently associated with plasmids (10%) than with chromosomes (7%) in *K. pneumoniae*.

In *A. baumannii*, 97 AmpC were detected in 122 chromosomal genome assemblies. Furthermore, 327 AmpC were found on chromosomes, and 1 on plasmids in 457 genomes containing both chromosomes and plasmids, indicating a higher prevalence of AmpC on chromosomes (16%) compared to plasmids (0.1%) and more than one copy of AmpC might be present on chromosomes of *A. baumannii* strains.

For *P. aeruginosa*, 250 AmpC was observed in 497 complete chromosomal genome assemblies, whereas 75 AmpC was found on chromosomes and 1 AmpC on plasmids across 127 genomes containing both chromosomes and plasmids. This indicates that AmpC is predominantly localized on chromosomes (61%) rather than plasmids (0.8%) in *P. aeruginosa*.

*Enterobacter spp*. revealed the presence of AmpC in 19 out of 98 chromosomal genomes. In addition, 411 genomes with both chromosomes and plasmids showed 233 AmpC on the chromosomes and none on the plasmids. This clearly showed that AmpC was more prevalent on chromosomes (10%) than on plasmids (0%) in *Enterobacter spp.* Based on our observations, we may conclude that the distribution and localization of AmpC varied across ESKAPE pathogens. AmpC was absent in the *E. faecium* and *S. aureus* genomes but present in *Enterobacter spp*. genomes in small amounts. It was highly prevalent on chromosomes in *A. baumannii* (∼1.7 copies per genome) and *P. aeruginosa* (∼1.3 copies per genome), followed by *K. pneumoniae* and *Enterobacter spp*. AmpC was the most common ESKAPE pathogen, with around 1.5 copies per genome in *A. baumanni*i, followed by *P. aeruginosa* and *K. pneumoniae.* Our results are aligned with a recent study that highlighted the significant chromosomal localization of AmpC in *A. baumannii* and *P. aeruginosa* and highlighted the possible contribution of these pathogens to antibiotic resistance [16, 17, 18].

### 2.4 Classification of *blaAmpC* into different enzyme groups

1790 AmpC proteins in this study formed nine clusters; hence, the *blaAmpC* discerned in the ESKAPE pathogens were further classified into nine enzyme groups based on their sequence homology and information provided in the BLDB database [15]. The comprehensive details about the identified enzyme groups, related variations, and the organisms with their gene location that contain these enzyme groups among the ESKAPE pathogens are provided in (**Table 2 & S1)**.

**Table 2:** List of variations within each enzyme group by comparing amino acid positions where substitutions occur.

| S. No. | Enzyme Group | Organism & Gene Location (s) | Number of sequences | Variations |
| --- | --- | --- | --- | --- |
| 1 | CMY | *K. pneumoniae* (B: chromosome, B: plasmid, O: chromosome), *P. aeruginosa* (B: Plasmid) | 54 | A171S, R221L, R221W, N366I, W374R |
| 2 | DHA | *K. pneumoniae* (B: plasmid, B: chromosome, O: chromosome)<br>*Enterobacter spp* (B: chromosome) | 135 | D165Y |
| 3 | ADC | *A. baumannii* (B: chromosome, B: plasmid, O: chromosome) | 755 | R2Q, L11F, S12P, S18N, G24D, D29E, I32V, E45D, M52I, V56I, M65I, S80R, S81I, T82S, T94N, A95G, G99A, N103A, D110K, T112R, T112K, E119V, P124S, D126N, G139S, Q144K, Q144R, K150Q, D152E, F158L, K163Q, P164T, S167A, S167P, Y171S, R172Q, G183R, K184E, V185I, V185F, A187G, A187S, N191K, K192Q, F194L, D195S, V197A, K200E, A205D, A205P, G207H, H210N, H210Y, P216A, M220I, A224D, F225Y, F225V, V236A, N237T, P238R, D242G, P254A, A255E, G257S, S267K, S267T, H270Y, H270N, L273F, N274T, P279S, A280K, A280T, D281N, N287S, R293H, R293F, V296E, V296I, V296L, N297G, S309A, A312T, T313P, T313M, N321S, T333S, Y344F, T351N, T351S, G352R, T355S, |
|  |  |  |  | V357I, A386T, N389D,<br>N389S, K393E |
| 4 | ACC | *Enterobacter spp* (B:<br>chromosome) | 13 | Not found |
| 5 | ACT | *Enterobacter spp* (B:<br>chromosome) | 377 | M36K, K37T, K37R, S39F,<br>S39L, S39T, L40V, L40F,<br>C41F, C42S, L45M, G47S,<br>G47A, I48L, I48V, I48T,<br>C50G, S51A, A52V,<br>A55G, A55S, A55T, P56S,<br>V57L, V57M, Q61E,<br>A63S, E64K, E64D,<br>V65M, A67K, A67E,<br>N68Q, N68R, T69N, V70I,<br>K75I, S78G, S78A, S78V,<br>V79I, Q89E, K91H, K91Q,<br>P92S, Y95F, K99E,<br>V102T, I102V, A103T,<br>A104K, N105S, N105K,<br>K106T, P107S, I119V,<br>G125A, I132V, D140G,<br>D140N, P142A, T144I,<br>T144A, R145K, Q149D,<br>Q149E, Q149A, R159S,<br>M160L, D177E, E178N,<br>E178D, V179I, T180K,<br>T180A, N182D, N182P,<br>A183T, S184A, R187N,<br>N191D, N191H, N191Y,<br>N191T, T200A, A207T,<br>A207S, K218Q, P223R,<br>P223N, P223G, P223S,<br>Y224F, Q226R, A227T,<br>M228I, T230K, T230E,<br>L233F, P235R, K237R,<br>K237A, D239N, D239H, |
|  |  |  |  | N244Q, N244D, A248S,<br>A248E, E249D, E250A,<br>A251T, A251Q, H252Y,<br>W255R, A262T, V263I,<br>V263T, R264H, M269T,<br>A272T, A272G, Q273E,<br>T279S, V281I, V281A,<br>Q282E, Q282K, M284I,<br>A285S, N286S, V288L,<br>M289K, A290V, A293N,<br>A293V, A293K, E295D,<br>E295A, N296A, N296K,<br>N296R, N296S, V297L,<br>A298E, A298Q, A298S,<br>A298P, D299N, D299A,<br>A300N, A300S, A300T,<br>S301T, K303G, K303R,<br>Q304K, Q304E, I306L,<br>A307S, A307T, I316V,<br>S318A, E333D, A334G,<br>A334S, N335Q, N335K,<br>V337I, V338I, E339D,<br>E339G, S343N, V345I,<br>P349A, P351L, V352A,<br>A353R, A353V, N356S,<br>A359V, K363E, K363N,<br>T370M, Q387E, Q387N,<br>I388L, I388V, K396T,<br>S397N, H409Q, H409R,<br>E412S, E412N, E412D,<br>E412K |
| 6 | CMH | *Enterobacter spp* (B:<br>chromosome,<br>B: plasmid, O: chromosome) | 16 | E25K, G123S, D144N,<br>A149V, P217S, H218N,<br>P264S, Q265P, A300V,<br>D378E, A379T |
| 7 | MIR | *Enterobacter spp* (B:<br>chromosome,<br>B: plasmid, O: chromosome) | 40 | A15T, S17A, K26T,<br>M40I, N41K, I103V,<br>A104S, A110T, R125H,<br>D128E, T148N, V198I,<br>K203N, D205N, A214T, |
|  |  |  |  | E225D, K248Q, V254L, I255M, A256V, Q264H, A265D, P266S, K269R, Q270K, R281C, G305E, V318A, A319R, P327L, K329N |
| 8 | PDC | *P. aeruginosa* (B: chromosome, B: plasmid, O: chromosome) | 397 | D3H, T4A, R5T, R5G, F6I, P7L, P7S, C8N, I12L, L18F, F19L, T21A, P23S, I25F, G27D, G27T, D32E, A36T, A55T, S59T, L71V, G77A, R79Q, R79K, P82L, T94S, A97V, A105T, R114S, Q117L, Q117R, A136G, F147L, S150V, Q155R, Q155P, A156V, A156T, A170T, Q174H, L176Q, L176R, G183D, E198K, L200S, L200I, Q203R, Q204R, V205L, Q213H, H215F, L216I, D217R, E220T, A222Q, L223Q, A224G, Q225L, K232N, R235H, V239A, E247G, G248A, V251L, T253S, D263E, D272E, R273K, P274T, P274Q, P274L, S306A, S306T, S306D, L310M, T316A, P322A, R324K, I325V, A330V, E335D, V353L, V356I, L361V, I365V, G391P, G391A, K392N, K396V, K396A |
| 9 | PIB | *P. aeruginosa* (O: chromosome) | 3 | Not found |

The maximum number of *blaAmpC* sequences belong to the ADC enzyme group (755 proteins), followed by the PDC enzyme group (397 proteins), ACT enzyme group (377 proteins), DHA enzyme group (135 proteins), CMY enzyme group (54 proteins), MIR enzyme group (40 proteins), CMH enzyme group (16 proteins), ACC enzyme group (13 proteins), and PIB enzyme group (3 proteins). Interestingly, the bigger enzyme group, ADC, was exclusively present in both plasmids and chromosomes of *A. baumannii* strains (**Table 2**). Therefore, ADC enzymes are chromosomally encoded and appear to be an imprint inherent to *A. baumannii*. The ADC β-lactamase is regarded as “universally present in the *A. baumannii* chromosome” [19], emphasizing how widespread and conserved it is within this particular species. This is consistent with the earlier study’s finding of the complete presence of ADC enzyme group in A. baumannii. [20] further supports this, reporting a positive rate of 72% for the *blaAmpC* in *A. baumannii* isolates, which all contained ADC-type AmpC resistance genes. The ACT enzyme group (377 proteins), CMH enzyme group (16 proteins), MIR enzyme group (40 proteins), and ACC enzyme group (13 proteins) were present exclusively in *Enterobacter spp.,* seen in previous studies also [21, 22, 23, 24]. PDC enzyme group (397 proteins) and PIB enzyme group (3 proteins) were exclusively present in *P. aeruginosa*. This was also reported in the previous studies [25, 26]. The remaining two enzyme groups, such as CMY (54 proteins), were present in *K. pneumoniae* and *P. aeruginosa* [27, 28], while DHA (135 proteins) were present in *K. pneumoniae* and *Enterobacter spp.* [29, 30].

### 2.5 Evolutionary relatedness among *blaAmpC* enzyme groups

Phylogenetic analysis of the 1790 *blaAmpC* sequences revealed that AmpC enzyme groups of *A. baumannii, P. aeruginosa, K. pneumoniae, and Enterobacter spp.* formed four distinct groups, each suggesting conserved evolutionary paths within our groups (**Figure 1**). The AmpC sequences of *P. aeruginosa* and *A. baumannii* formed distinct and well-defined groupings, indicating significant conservation. *K. pneumoniae*, on the other hand, had mixed clustering with *P. aeruginosa* and *Enterobacter species*. Additionally, *Enterobacter spp.* and *P. aeruginosa* had mixed clustering, suggesting that their *blaAmpC* share evolutionary origins unique to these genera. The phylogram of AmpC enzyme groups further highlights that these evolutionary patterns are independent of the genetic location (chromosomal/plasmid) among all the AmpC enzyme groups **(Figure S1)**. AmpC enzyme groups ACC, ACT, ADC, CMH, CMY, DHA, MIR, PDC, and PIB, which were present in all the four pathogens, formed separate clusters, but these clusters could be sub-grouped based on the pathogen. However, AmpC enzyme groups ADC, ACC, PDC, ACT, MIR, CMH, and PIB are present only in a single pathogen and form independent clusters, respectively.

**Figure 1:**
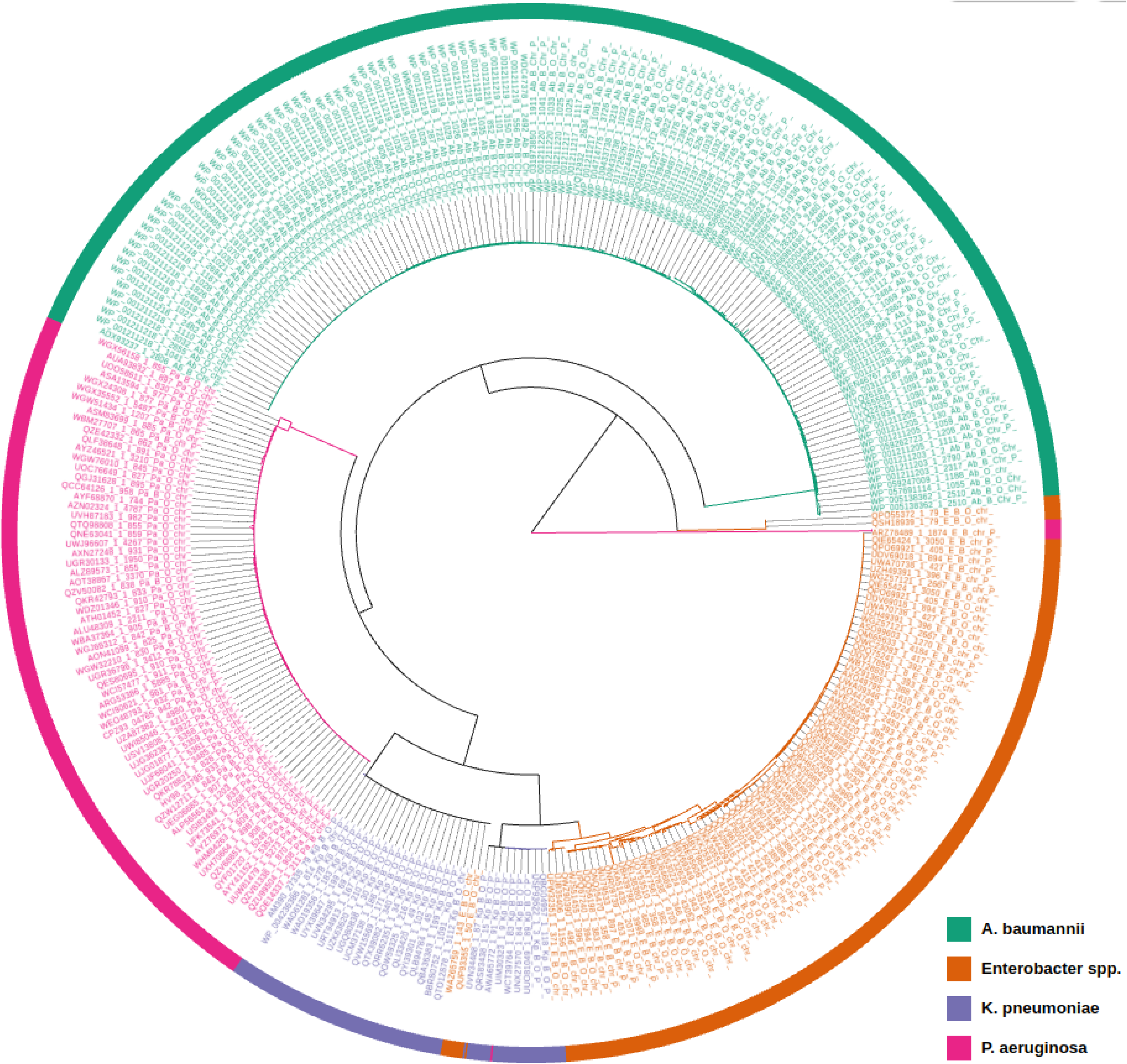
A phylogram of 1790 AmpC sequences from *A. baumannii, Enterobacter spp., K. pneumoniae, and P. aeruginosa*, illustrating the evolutionary relationships and diversity among ESKAPE pathogens. The tree session is also available for broader exploration at this link: https://microreact.org/project/v9BTczaE4WQGH5AvxB6CQN-pathogencirc. In this shared session, users can zoom in and out, and scroll (up, down, left, and right) to examine specific branches and clades of each enzyme group and their corresponding sequences.

### 2.6 Phylogenetic insights into *blaAmpC* enzyme groups

Amino acid variation analysis within the nine AmpC enzyme groups revealed that the amino acid sequences of the ADC enzyme group formed a separate cluster, which is exclusively conserved in *A. baumannii*. The PDC conserved in *P. aeruginosa* forms its cluster. Since DHA is present in both *Enterobacter spp.* and *K. pneumoniae*, it is possible that these pathogens shared a genetic ancestor or an evolutionary pathway. *P. aeruginosa* and *K. pneumoniae* share the CMY enzyme group, which may indicate horizontal gene transfer. Enzyme groups restricted to *Enterobacter spp*. form unique clusters, like ACC, ACT, MIR, and CMH, which show diversification within their enzyme groups. PIB is present exclusively in *P. aeruginosa*, forming a unique branch with a branch length of 3.3812, which likely reflects its significant evolutionary rate and divergence from the other eight *blaAmpC* enzyme groups. Notably, the largest enzyme group, ADC, showed the most significant degree of variation, followed by ACT and PDC, indicating a notable degree of diversity within these groups (**Table 2**). On the other hand, the ACC and PIB enzyme groups in *Enterobacter spp.* showed no variations, indicating potential conservation of function within these enzyme groups (**Table 2**).

### 2.7 Pairwise identity between the enzyme groups

Given the diversity of *blaAmpC* amongst different pathogens; we conducted pairwise comparisons of the representative sequences of nine enzyme groups to examine the relatedness among them. The pairwise identity between the entities ranged from 18.84% to 100% (**Table 3**). For instance, the PIB and ACC pair showed only 18.84% identity, whereas the ACT and MIR pair showed 89.76%. The other AmpC enzyme groups, such as CMH and MIR, showed 89.50%, CMH and ACT showed 87.40% and CMH and CMY showed 76.38% identity, respectively, which indicates that these entities are closely related. However, PIB and ADC & ACC, ADC, and CMY had a lower identity of 18.84% and 35.47 respectively, indicating a higher degree of divergence or they might be distant homologs. These findings demonstrate the diversity through percentage identity among the various entities, with some pairs showing close alignments and others showing significant divergence.

**Table 3.** Pairwise amino acid identities (%) between AmpC enzyme groups.

|  | PIB | ADC | ACC | PDC | DHA | CMY | CMH | ACT | MIR |
| --- | --- | --- | --- | --- | --- | --- | --- | --- | --- |
| PIB | 100.00 |  |  |  |  |  |  |  |  |
| ADC | 20.47 | 100.00 |  |  |  |  |  |  |  |
| ACC | 18.84 | 37.11 | 100.00 |  |  |  |  |  |  |
| PDC | 18.84 | 39.95 | 42.64 | 100.00 |  |  |  |  |  |
| DHA | 20.66 | 38.61 | 36.97 | 46.56 | 100.00 |  |  |  |  |
| CMY | 20.66 | 35.47 | 39.42 | 41.84 | 56.20 | 100.00 |  |  |  |
| CMH | 20.96 | 39.20 | 38.89 | 45.00 | 55.41 | 76.38 | 100.00 |  |  |
| ACT | 19.76 | 38.13 | 38.10 | 44.47 | 53.30 | 74.02 | 87.40 | 100.00 |  |
| MIR | 20.36 | 37.60 | 38.10 | 43.95 | 55.67 | 75.85 | 89.50 | 89.76 | 100.00 |

### 2.8. Amino acid analysis of representatives of the AmpC enzyme groups and their phylogenetic study

The AmpC enzymes are a heterogeneous group, with diverse sequences and functional properties. All enzyme groups of *blaAmpC* belong to Ambler class C, however, we found low sequence identity within the class. This means that although these enzymes have similar functions (i.e., β-lactamase activity), their amino acid sequences differ significantly. Hence, to better understand the common characteristics that might be shared among the various *blaAmpC* enzyme groups, the amino acid sequences of representative sequences of the nine *blaAmpC* enzyme groups were analyzed. The analysis aims to identify conserved motifs, structural features, or functional domains that explain enzyme properties like substrate specificity, stability, and antibiotic resistance profiles by aligning representative sequences.

Green bars in the identity plot of **Figure 2** highlight highly conserved regions observed across all the representative sequences; these sequences suggest functional and structural importance, as evolutionary pressures preserve these residues to maintain the essential role. Notably, all three significant conserved motifs, ‘SXSK’, ‘YXN’, and ‘KTG’ of class C β-lactamase were conserved in all eight enzyme groups excluding the PIB enzyme group. However, PIB retains the SXXK motif, which is essential for its catalytic activity, but lacks the typical YXN and KTG motifs that are present in other eight enzyme groups (e.g., ACT, MIR, CMY, DHA, PDC, ADC, ACC). Instead of the YXN and KTG motif, it possesses the YST and AQG, respectively [31].

**Figure 2:**
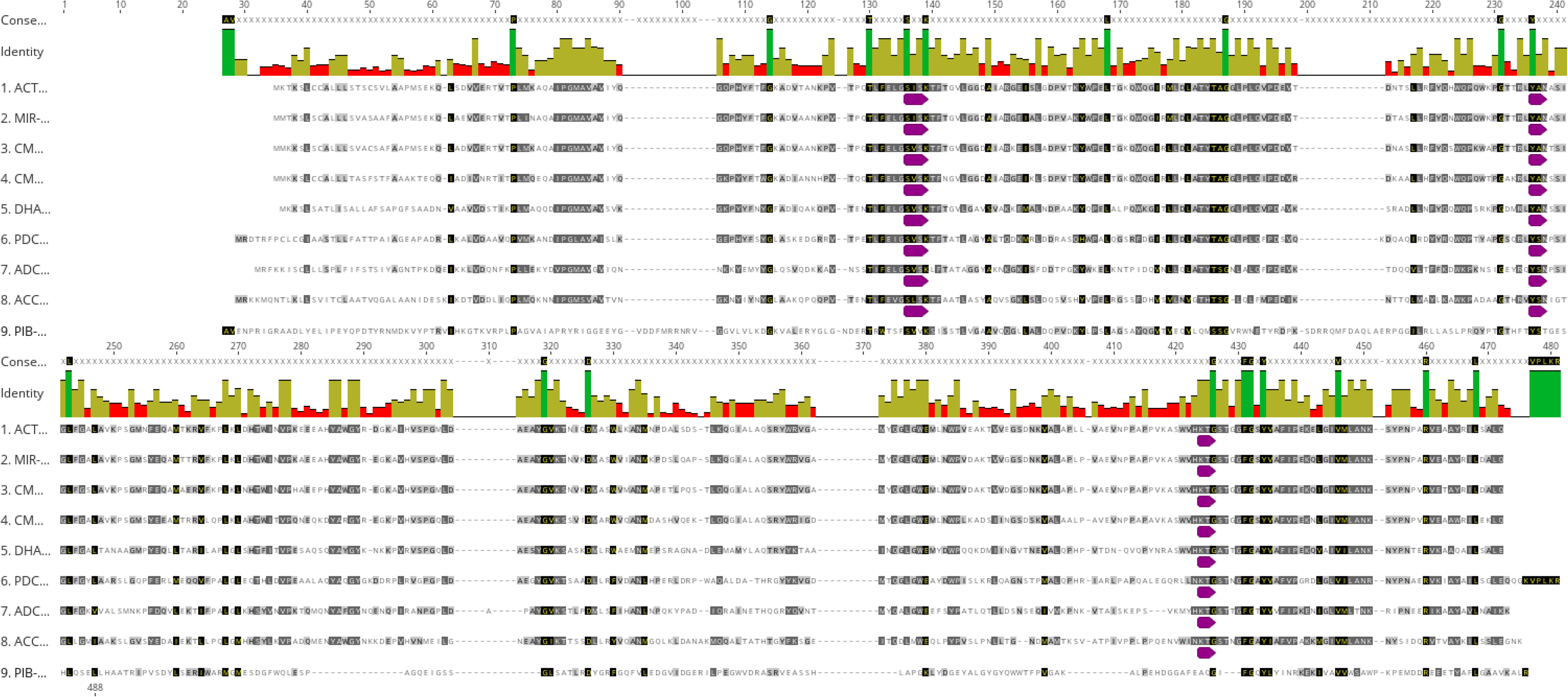
A multiple sequence alignment (MSA) of representative sequence from each of the nine *blaAmpC* enzyme groups. In the corresponding conservation plot, identical amino acid residues across the sequences are highlighted in dark green, providing a clear visual indication of conserved regions within the enzyme groups. This plot helps to elucidate conserved motifs and evolutionary patterns among the different *blaAmpC* enzymes.

A time-stamped phylogenetic tree constructed using nine representative sequences of AmpC enzyme groups formed two distinct main branches, evolved from common ancestors (**Figure 3**). The first branch forms an outgroup of the PIB enzyme group, underscoring the enzyme group’s uniqueness within the *blaAmpC*. Despite being categorized as an AmpC Class C β-lactamase, the enzyme group showed a completely different pattern from the other eight. This was also confirmed in the study of *Iorga et al.*, which indicates that PIB does not bind to cephalosporins while enhancing its activity against carbapenems [31] because of the difference in conserved motifs. PIB is excluded from our further study because we mainly focus on cephalosporins-based antibiotic resistance groups.

**Figure 3:**
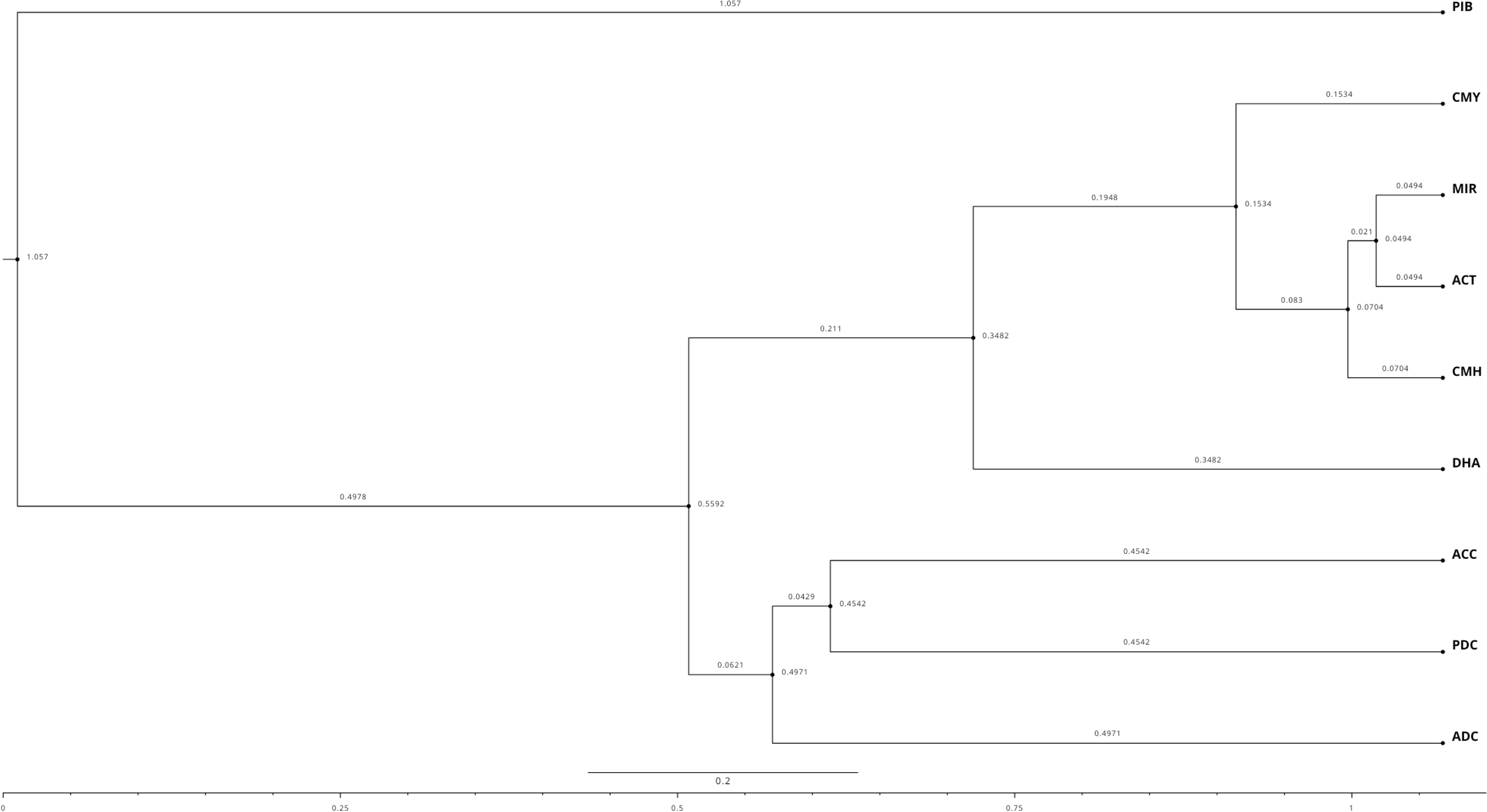
A timestamped phylogenetic tree of a representative sequence from each of the nine *blaAmpC* enzyme groups. Timestamp provides insights into the chronological progression and the emergence of distinct *blaAmpC* enzyme groups, illustrating the evolutionary relationships and divergence of the sequences over time.

However, the second main branch further splits into two sub-branches, which, with time, are divided into eight sub-branches representing eight different enzyme groups of *blaAmpC* (**Figure 3**). Further, ACT, MIR, CMY, and CMH enzyme groups belong to the same branch, hence, the members of these subfamily pairs might have evolved from the same ancestor. This is also supported by the pairwise identity between MIR and ACT, MIR and CMH, ACT and CMH, CMH and CMY, MIR and CMY, and ACT and CMY enzyme groups which were 89.76%, 89.50%, 87.40%, 76.38%, 75.85%, and 74.02% respectively. These enzyme groups’ branch lengths and the time elapsed since their divergence from a common ancestor, also demonstrated in their sequence identities. The sequence identities indicate a recent divergence or a close relationship between them. Overall, this suggested that higher the pairwise sequence identity between two sequences, the smaller the divergence between them and the shorter the period since their divergence.

### 2.9. Structural variations in 3D protein models of *blaAmpC*

The superimposition of 3D protein structures of the representatives of each AmpC enzyme group demonstrated that the protein structures were highly superimposable (**Figure 4**). The details of the Root Mean Square Deviation (RMSD) values of pairwise structure alignments of the enzyme groups are shown in **Table 4**. The maximum RMSD value was observed in ADC and MIR enzyme groups, which showed a pairwise sequence identity of 37.560%. In the phylogenetic tree, they also represent two separate branches and form different clades. Interestingly, the RMSD values of pairwise structural alignments of two AmpC enzyme groups, namely MIR and ACT; MIR and CMH; ACT and CMH; CMH and CMY; MIR and CMY, and, ACT and CMY enzyme groups showed minimum RMSD values in Å as 0.616, 0.8, 0.817, 1.056, 0.636 and 0.666, respectively. These enzyme group pairs showed 89.76%, 89.50%, 87.40%, 76.38%, 75.85%, and 74.02% identity, respectively. In the phylogenetic tree, CMY, MIR, ACT, and CMH have the same origin, also MIR and ACT have the same branches without any divergence. Consequently, it can be deduced that a low RMSD value in pairwise alignment correlates with a high pairwise sequence identity and a low degree of divergence. The multiple sequence alignment of the AmpC sequences demonstrated that the primary and secondary amino acid sequences were significantly conserved within a subfamily (**Figure S2**).

**Figure 4:**
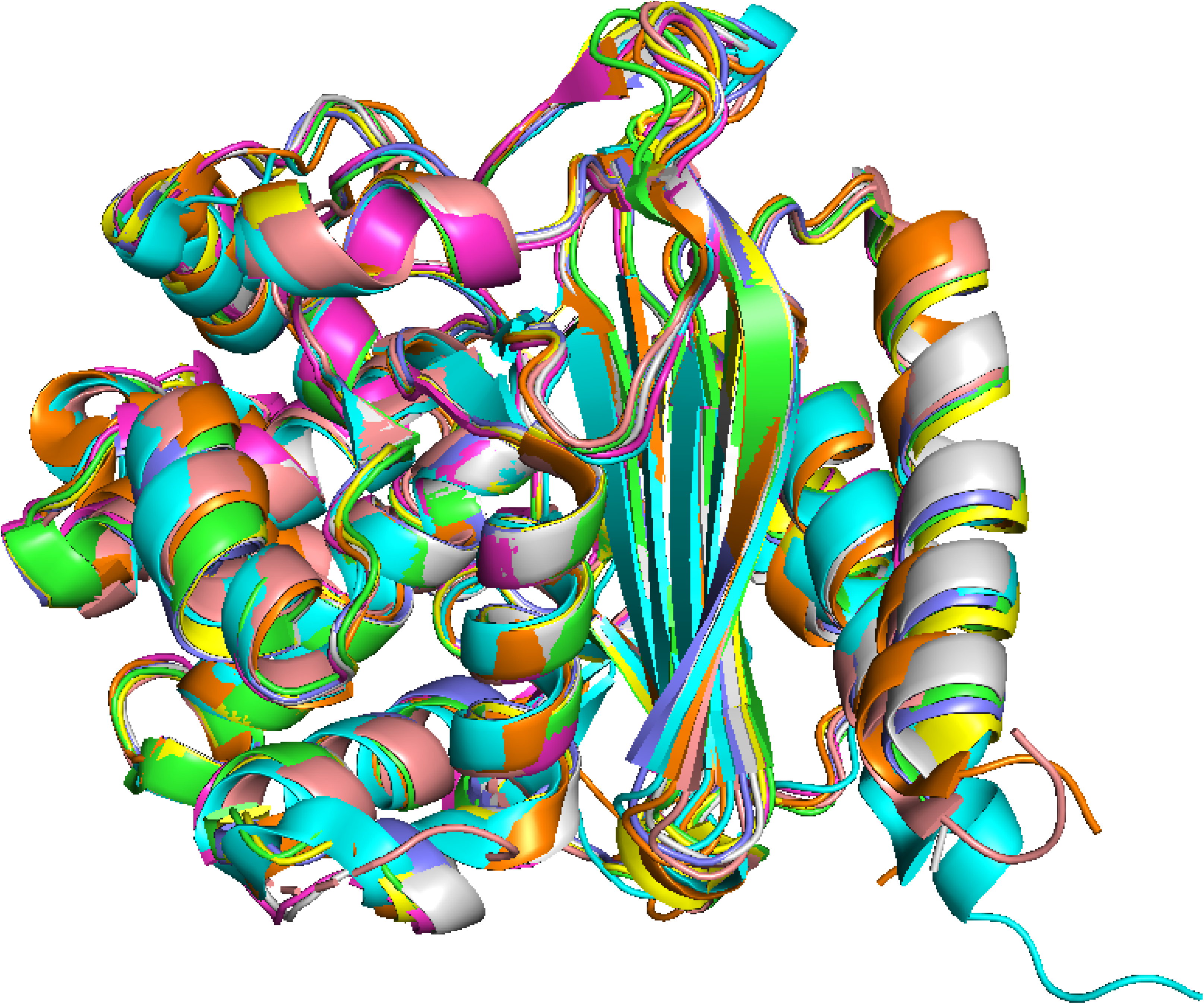
Superimposed 3D protein models of representatives of the nine blaAmpC enzyme groups. (Color Key, Orange: 8FQV(ADC), Light Pink: 6K8X(ACC), Cyan: 6S1S(PDC), Yellow: 5GGW(DHA), Green: 1ZC2(CMY), Grey: 6LC7(CMH), Purple: 7TI1(ACT), Magenta: 2ZC7(MIR).

**Table 4.** Root Mean Square Deviation values of pairwise structure alignments of 3D protein models of the nine AmpC enzyme groups (The color scale is from highest (green) to lowest (red))

|  | ADC | ACC | PDC | DHA | CMY | CMH | ACT | MIR |
| --- | --- | --- | --- | --- | --- | --- | --- | --- |
| ADC | 0 |  |  |  |  |  |  |  |
| ACC | 1.729 | 0 |  |  |  |  |  |  |
| PDC | 1.395 | 1.267 | 0 |  |  |  |  |  |
| DHA | 1.683 | 1.455 | 1.188 | 0 |  |  |  |  |
| CMY | 1.628 | 1.308 | 1.287 | 0.543 | 0 |  |  |  |
| CMH | 1.762 | 1.291 | 1.407 | 1.042 | 1.056 | 0 |  |  |
| **ACT** | 1.641 | 1.337 | 1.205 | 0.541 | 0.666 | 0.817 | 0 |  |
| **MIR** | 5.283 | 1.341 | 1.175 | 0.546 | 0.636 | 0.8 | 0.616 | 0 |

## 3. Materials and Methods

### 3.1. Data Collection

The genomes of *E. faecium, S. aureus, K. pneumoniae, A. baumannii, P. aeruginosa, Enterobacter spp*. (ESKAPE pathogens) were downloaded from the NCBI genome database (September 21, 2023). The number of *E. faecium, S. aureus, K. pneumoniae, A. baumannii, P. aeruginosa, and Enterobacter spp.* genomes in the database were 3460, 15,739, 17608, 8037, 8221, and 4970, respectively. Incomplete genome assemblies were explicitly excluded from further study to limit any potential exclusion of AmpC enzyme groups. **Table 1** provides an overview of the statistics throughout the data-gathering process. Depending on the nature of the replicons harbored in the genome, we classified each genome into two categories: (a) those with only chromosomes and (b) those with both chromosomes and plasmids. Afterward, each replicon was searched for the *blaAmpC*.

### 3.2 Searching of *blaAmpC* in ESKAPE sequences

The profile Hidden Markov Models (pHMM) of *blaAmpC* were retrieved from the β-LacFampred tool [13]. Each replicon was searched against the pHMM of AmpC using the NHMMSCAN program of the HMMER package (version 3.1) [14] with an E-value threshold of 1e-06. The NHMMSCAN search results were validated using the GenBank annotations of the corresponding genomes to retain only the verified *blaAmpC* and eliminate non-AmpC sequences. Afterwards, the translated protein sequences of verified *blaAmpC* were retrieved from the NCBI protein database.

### 3.3 Distribution of *blaAmpC* into enzyme groups

The protein sequences of class C β-lactamases were obtained from the comprehensive, manually curated public resource of β-lactamases known as BLDB (Beta-Lactamase DataBase) [15]. A comprehensive BLAST search identified *blaAmpC* sequences across the ESKAPE pathogens. Only the sequences with 100% identity were taken to ensure the reliability of the data. This refinement was done to avoid non-*blaAmpC* sequences.

### 3.4. Multiple sequence alignment and phylogenetic analysis

Multiple Sequence Alignment (MSA) was done using the MAFFT program [32] and visualized using Jalview [33]. A Neighbor-Joining (NJ) method followed by 1000 bootstrap values was used for robust tree construction. The phylogenetic tree was visualized using Microreact [34], a web-based application to investigate the evolutionary patterns of *blaAmpC* across the ESKAPE pathogens.

### 3.5. Assessment of pairwise sequence identity in AmpC enzyme group representatives and construction of bayesian time-stamped phylogenetic trees

A representative protein sequence from each AmpC enzyme group was used to compare amino acid sequences and construct the phylogenetic tree. We identified the representative sequences from the multiple sequence alignment (MSA) of each *blaAmpC* enzyme group. The first/topmost sequence of the MSA was considered the representative of that enzyme group. The pairwise sequence identity of the nine AmpC enzyme group representatives were calculated to determine the evolutionary relationships among them. Time-stamped phylogeny was constructed using Bayesian Evolutionary Analysis Sampling Trees (BEAST) [35], a cross-platform program for Bayesian evolutionary analysis, and MCMC sampling to infer the evolutionary patterns. Before using BEAST, alignment of the representative sequence was done and the tree file in nexus format was generated using ClustalW 2.1 program [36], which was further used for BEAST analysis. The nexus format tree file was used to load the parameters using the BEAUti program setting the substitution model and clock model to default, the parameters were saved in .xml format. BEAST was run on .xml, where MCMC is run for 10,000 iterations, saving samples every 1,000 iterations.

The BEAST output log file was analyzed using the TRACER program. TreeAnnotator was employed to condense the posterior sample into a maximum clade credibility (MCC) tree. The configuration included a Burn-in of 50, a Posterior Probability Limit of 0.0, and the selection of a maximum clade credibility tree with default settings. The mean heights option was selected for node heights, which assigns the average height across the entire tree sample to each node in the tree for that clade. Finally, the tree was visualized using FigTree, which showed node ages, branch lengths, scale bars, axis, and posterior probability for each node.

### 3.6. 3D structural variations in the AmpC enzyme groups

The 3D structures of representative sequences of each AmpC were retrieved through BLAST search against the PDB. The AmpC protein structures obtained using BLAST search were selected based on % alignment identity and % query coverage. For eight enzyme groups, namely ADC, ACC, PDC, DHA, CMY, CMH, ACT and PIR protein structures were found in the PDB database, which showed more than 60% identity and 92% query coverage. Hence, these structures were used as representatives of their AmpC enzyme groups. The protein name, PDB ID, query coverage, and group members’ identities are shown in **Table 5**. Further, all eight protein structures were aligned and assessed by superimposing them using PyMOL Schrodinger. The RMSD values of all 3D models were then calculated for pairwise structural comparison. **Figure 5** depicts the overall workflow adopted in this study.

**Figure 5:**
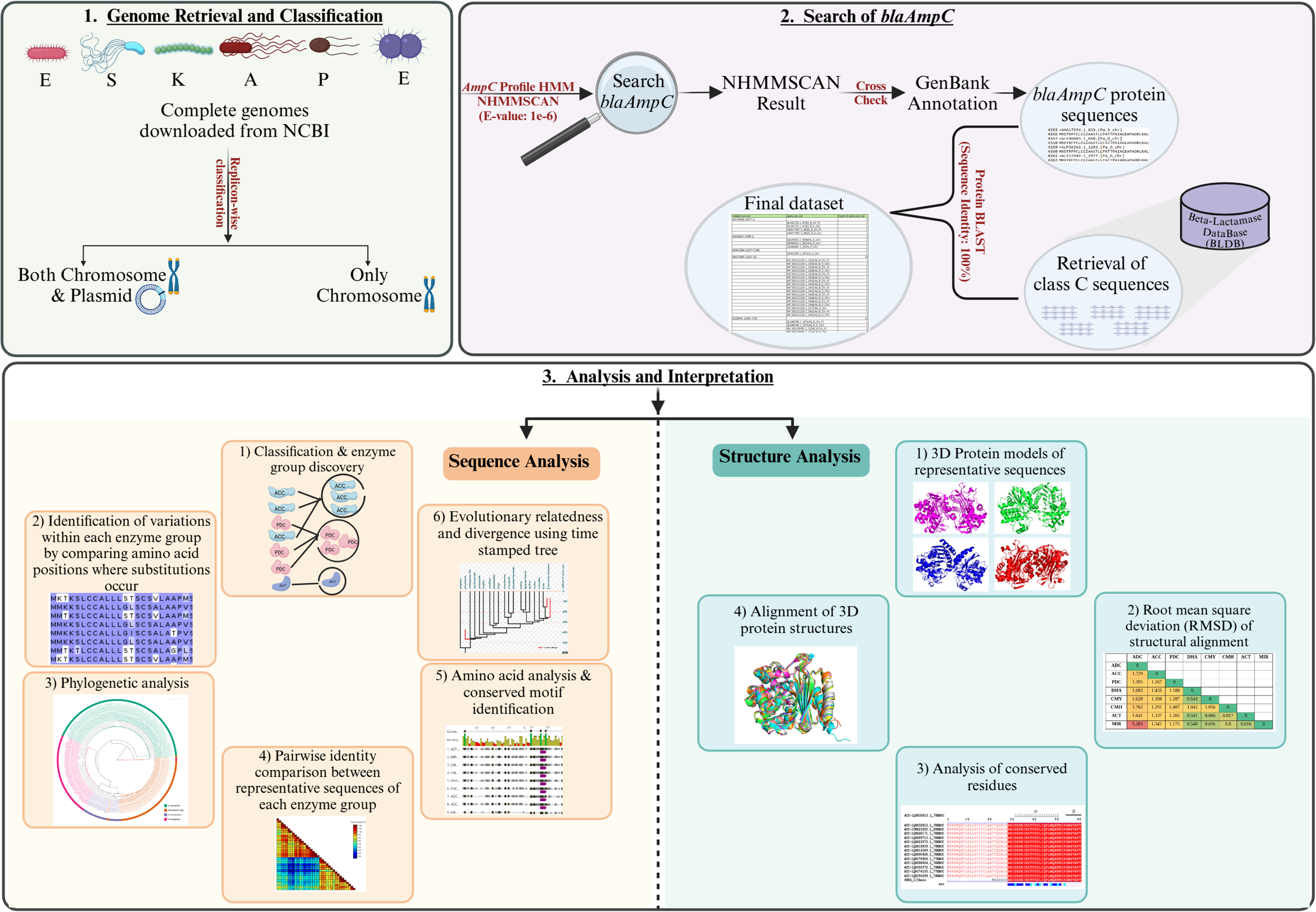
The comprehensive methodological approach employed for the investigation of *blaAmpC* in ESKAPE pathogens.

**Table 5.** PDB IDs of the structural representatives of the nine AmpC enzyme groups with query coverage and identity percentage.

| S. No. | Protein Name | PDB ID | Query Coverage (%) | Identity (%) |
| --- | --- | --- | --- | --- |
| 1 | ADC | 8FQV | 93% | 99.72% |
| 2 | ACC | 6K8X | 93% | 100.00% |
| 3 | PDC | 6S1S | 100% | 100.00% |
| 4 | DHA | 5GGW | 92% | 61.25% |
| 5 | CMY | 1ZC2 | 94% | 99.72% |
| 6 | CMH | 6LC7 | 94% | 97.51% |
| 7 | ACT | 7TI1 | 100% | 86.09% |
| 8 | MIR | 2ZC7 | 94% | 93.59% |

## 4. Conclusion

This study provides a detailed overview of *blaAmpC*, one of the most important AMR enzymes, in ESKAPE pathogens to study their occurrence, localization, classification, and detailing the molecular, evolutionary, and structural characteristics. Through the analysis of 4713 complete genomes of ESKAPE pathogens, 1790 distinct AmpC enzymes were found, revealing the absence of AmpC in *E. faecium* and *S. aureus*. Nine enzyme groups were identified on the grouping of the enzymes: ADC, PDC, ACT, DHA, CMY, MIR, CMH, ACC, and PIB. Our study revealed the exclusive presence of ADC in *A. baumannii*, PDC, and PIB in *P. aeruginosa*, and ACC, ACT, CMH, and MIR in *Enterobacter spp*. Variations within individual enzyme group sequences (intra-group) were identified by comparing amino acid sequences where substitution occurred, which revealed that the enzyme group ADC exhibited the greatest sequence diversity, followed by PDC and ACT. Phylogenetic analysis highlights the recent divergence and close relationships between enzyme groups, revealing shared evolutionary origins or horizontal gene transfer. Amino acid sequence analysis highlighted conserved motifs like “SXSK,” “YXN,” and “KTG,” which are necessary for β-lactamase activity, which was absent or altered in the case of the “PIB” enzyme group, modifying its functional properties, reducing its affinity for cephalosporins and increasing its activity against carbapenems. A similar pattern was observed in the timestamp phylogenetic tree where “PIB” formed an outgroup. This divergence made us exclude “PIB” from further analysis, focusing on cephalosporins. Structural analysis further supported the evolutionary relationships observed in phylogenetic analysis, found through the superimposition of 3D structures, where lower RMSD values were associated with higher sequence identity and lower divergence. Our results highlight *blaAmpC* enzymes contributing to the escalating problem of antimicrobial resistance among strains in the nosocomial environment. Therefore, we emphasize the responsible use of antibiotics, particularly in hospital settings such as Intensive Care Units (ICUs) that raise serious concerns about limited treatment options, to ensure more effective treatments and better patient outcomes in the fight against AMR.

## Acknowledgements

All authors thank the International Centre for Genetic Engineering and Biotechnology, New Delhi (India) for providing facilities to pursue the research work.

## Author’s Contributions

DP conceived the idea, collected and organized the data. DP and IG performed the analysis, finalized the results and drafted the original draft. DG supervised the work and revised the manuscript. All authors reviewed and finalized the manuscript.

## Data availability

Not applicable

## Declarations

### Ethics approval and consent to participate

Not applicable

### Consent for publication

Not applicable

---

**Figure S1:** A phylogram of 1790 AmpC sequences, with each enzyme group represented by a distinct color shade. The phylogenetic tree can be explored in more detail through the following shared session: https://microreact.org/project/okssuk9PwZ7tUmZwsMp2f6-familycirc. The session allows for interactive zooming and scrolling (up, down, left, and right) to facilitate the examination of specific branches and clades of each enzyme group and their corresponding sequences.

**Figure S2:** Structure based multiple sequence alignment of the AmpC enzyme groups in ESKAPE Pathogens

